# Field decomposition of Bt-506 corn leaf and its effect on Collembola in the black soil region of Northeast China

**DOI:** 10.1101/792309

**Authors:** Baifeng Wang, Fengci Wu, Junqi Yin, Ling Zhang, Xinyuan Song

## Abstract

The litters of Bt corn would go into the soil through straw returning and field ploughing after cultivation. To clarify whether the leaf litter decomposition rate and the non-target soil Collembola were influenced by the Bt protein or other litter properties in leaf litters of Bt corn in Northeast China, leaf litterbags of Bt-506, its near isoline Zheng 58 and a local type Zhengdan 958 were used in the field in Northeast China. The leaf decomposition rate, the leaf properties and the collembolan community in litterbags were investigated later. After seven months, only 43.5 ng/g Bt protein in Bt-506 leaf litter was left. All the investigated indices were not significantly different between Bt-506 and its near isoline Zheng 58. But when compared with local type Zhengdan 958, Bt-506 and its near isoline Zheng 58 contained lower non-structural carbohydrate content but higher total nitrogen content, and had lower litter decomposition rate and less abundance of Collembola. Collembolan abundance and litter decomposition rate were both significantly correlated with the non-structural carbohydrate and total nitrogen contents of the leaf litters. Field study results revealed Bt protein did not affect the leaf litter decomposition rate and the collembolan community in leaf litterbags in short term. The significant differences of these investigated indices among corn types were caused by the different non-structural carbohydrate and total nitrogen contents in leaf litter.

## Introduction

Bt corn is developed to combat lepidopteran pests [1], and the wide planting of Bt corn can greatly reduce insecticide applications and labour requirements [2, 3]. The litters of Bt corns would go into the soil by straw returning and field ploughing. The Bt protein from these litters will persist for a long time, even more than 200 days in the soil [4].

The potential impacts of the Bt protein from transgenic crop on litter decomposition and non-target soil fauna have been reported in many studies. The results of the studies varied according to study materials and site. Most of the studies indicated that the Bt protein itself did not affect the decomposition rate of transgenic plant and non-target soil fauna [5-8]. However there were a few studies indicated that the decomposition rate of the litters and non-target soil fauna in the litters were directly related to the Bt protein for Bt crops [9, 10]. Besides, the litter property of Bt crop might have been changed by the transformation of exogenous gene. For example, Saxena and Stotzky [11] found that Bt corn had a higher lignin content compared with non-Bt corn, further Fang et al. [12] found that the lignin, lignin/nitrogen ratio and total organic carbon contents in Bt corn were higher than in non-Bt corn. Additionally, some researchers have found that the total amount of nitrogen per unit litter could influence fungal and bacterial populations in microcosms [13, 14]. Thus, the decomposition process of corn leaf litters [5, 12, 15], as well as non-target soil fauna in the field [16], were probably influenced by the significantly different leaf properties among the corn types through an influence on the activity of microbial communities.

Bt-506 is a new *Cry1Ac* corn in breed line that was independently developed by the China Agricultural University. In China, the black soil region is the most important location for corn production. Therefore, it is of great significance to assess the litter decomposition rule of Bt-506 corn and its effect on soil fauna in the black soil region of China.

Collembola are one of the most ubiquitous soil fauna, which include numerous species and play an important role in nutrient cycling of the soil system [17-19]. Generally Collembola can be good indicators of soil quality as they are sensitive to soil environmental change [20-22]. Collembola may directly or indirectly come into contact with the Bt toxin by feeding on the living root tissues and the litters of Bt plants, or soil fungi, microorganisms, and organisms (worms, protozoans) in the field with Bt plants [23, 24]. Some studies had been conducted to explore the effects of Bt plant cultivation on soil Collembola in recent years [5, 9, 25-28], while most of these studies were conducted during planting seasons [25-28]. At present, there were two studies were carried out in field to evaluate the effect of Bt litters on Collembola community at family level [5, 9]. The results didn’t show any major changes in the composition of the soil collembolan community. But one of the two studies [9] found that the species of the Tullbergiidae of Collembolan were much less in Bt crop filed than in non-Bt crop field. Furthermore, the two studies just identified the Collembola at family level. However, the Collembola is rich in species, and nearly 9000 species have been reported worldwide [29]. After a long term adaptation and evolution, the ecological niche and function of different collembolan species in the same family are not exactly the same [17, 18]. Thus investigating collembolan at species level is a more accurate way to study the response of collembolan community to environmental changes [25] and allows to analyse the response mechanism [28]. Till now, the effects of Bt crop litter on Collembola at species level were just carried out in laboratory conditions [30-34].

Therefore, we carried the following study in field conditions to assess the effect of Bt corn litter on soil Collembola at species level. Field trials were carried out in the black soil region of Northeast China in 2013 and 2014, with crop Bt-506, its near isoline Zheng 58, and a local crop type Zhengdan 958 as experimental materials. The litterbags of the three corn types were buried into the field after the autumn harvest, and then the remained litter weight, the Bt protein content, the non-structural carbohydrate content, the total nitrogen content and the collembolan community in the litter were investigated at three stages in the next year. The specific objectives of the present study were to clarify: 1) whether the transformation of exogenous gene affects the decomposition rate and leaf litter properties of Bt-506 corn in field; 2) whether the decomposition of transgenic Bt-506 corn affects soil Collembola at the species level in litterbag.

## Materials and methods

### Corn types and study field

Three corn types, transgenic corn Bt-506, its corresponding non-transformed near isoline Zheng 58 (control) and a predominant local corn type Zhengdan 958 (Zheng 58 ×Chang 7-2) were used in this study. Bt-506 is a newly developed *Cry1Ac* corn inbreed line by China Agricultural University. The seeds of Bt-506 and Zheng 58 were provided by China Agricultural University, and the seeds of Zhengdan 958 were provided by the Jilin Academy of Agriculture Sciences.

Corn were cultivated at an experimental farm of Jilin Academy of Agriculture Sciences in Gongzhuling City, China (43°19’ N, 124°29’ E). The annual average air temperature in experimental location was 5.6°C. The soil in this area was the typical black soil of Northeast China. Its characteristics are summarised as follows: pH, 6.31±0.03; organic matter, 27.08±0.07 g/kg; total nitrogen, 0.77±0.07 g/kg; available phosphorus, 10.68±0.07 mg/kg; and available potassium, 154.10±0.76 mg/kg.

A randomized block design involving the three corn types were established with three replications in 2013 and 2014. Blocks of the same corn type were not adjacent, and each type of corn was planted in the same blocks in 2014 as in 2013. Each block was 10 m by 15 m in size. Blocks were separated by a 2-m wide clearing. Corn plants were sown on May 25 in 2013 and on May 8 in 2014, at a density of 50,000 plants per ha. Field was cultivated using standardized agricultural management practices, and no insecticides were applied during the study. The corn plants were taken away on October 30, 2013, and then no cultivation practices was conducted until May 5, 2014.

### Leaves and leaf litterbags

On October 17, 2013 when corn matured, we randomly selected 10 plants from each corn type to collect senescent leaves without petiole. For each corn type, the leaf samples were put together. These corn leaves were cut into 5-cm pieces, dried at 40° for 72 h, and stored in a freezer at −20°C until use.

The litterbags (15 cm×10 cm) were made of nylon mesh with mesh size of 5 mm. Before put into litterbags, the leaf litters were dried at 40°C for 24 h. The 27 litterbags per corn type were prepared, 10 g per bag. On November 15, 2013, all litterbags were buried into the ridge soil at a depth of 5 cm. Each type of leaf litter was buried into the block where the same corn type was planted, and there were 9 litterbags in each block. To avoid edge effect, all the litterbags were buried more than 2 m away from the block border, and kept more than 1 m distance between every two adjacent litter bags. Plastic tag was set beside each bag. And to avoid destroying these litterbags and influencing the Collembola community in the litterbags, corn seeds were sowed more than 10 cm away from these litterbags in 2014.

### Sampling and analysis

The litterbags were taken out from soil on April 20, May 20 and June 20 2014, respectively, and for each time, three litterbags per corn type were sampled. Collembola in the litterbags were extracted using the Macfadyen method [35]. The extracted organisms were preserved in 95% ethanol for identification. Then the Collembola were identified at species or morphospecies level according to references [36-39].

After Collembola extraction, the leaf litters were poured out from the litterbags to the glass petri dishes, the soil mixed into the litters was took away carefully, then the leaf litters were dried at 40°C for 24 hours again, the leaf litters were weighed, the decomposition rates of the leaf litters were calculated. In addition, the contents of non-structural carbohydrate [40] and total nitrogen (Foss automated KjeltecTM instruments, Model 2100) of leaf litter were also analyzed. Bt protein content of Bt-506 leaf litter samples were quantified by enzyme-linked immunosorbent assay (ELISA) using a QuanliPlate Kit for Cry1Ab/Cry1Ac (Envirologix Inc., Portland, ME, USA).

### Data analyses

One-way ANOVA was performed using SPSS (version 23, IBM, USA) to analyze leaf litter decomposition rate, non-structural carbohydrate content, and total nitrogen content, corn type used as independent factor. A repeated measure ANOVA was performed to analyze the abundance and Shannon-Wiener index of Collembola species as well as the abundance of the dominant species *Proisotoma minuta*. In the analysis of repeated measure ANOVA, the sampling time was included as repetition levels, and the corn type was used as the “between subject” factor, and the number of collembolan individuals (*n*) was transformed by log_10_ (*<u>n</u>* + 1) to obtain a normal distribution. The person correlation analysis was also performed using SPSS (version 23) to analyze the relationships among collembolan abundance, leaf litter decomposition rate, non-structural carbohydrate content, and total nitrogen content.

## Results

### Bt protein, non-structural carbohydrate and total nitrogen contents

In the litterbags of Bt-506, the content of Bt protein in leaf litters was significantly decreased over time. The original content was 536.4 ng/g before it was buried into the field, it decreased to 420.9 in April 20, to 179.5 ng/g in May 20, and only 43.5 ng/g was left June 20.

Repeated measure ANOVA showed that the non-structural carbohydrate content of leaf litter was significantly higher (Table 1, *P* = 0.015), while the total nitrogen content was significantly lower (Table 1, *P* < 0.001) in local type Zhengdan 958 than in Bt-506 and its near isoline Zheng 58 types. As for each sampling time, one-way ANOVA analysis also showed that the non-structural carbohydrate content was significantly higher but the total nitrogen content was significantly lower in Zhengdan 958 than in Bt-506 and Zheng 58 types on April 20 and May 20 (Table 1, *P* < 0.05). And there was no significant difference in the non-structural carbohydrate and total nitrogen contents of three corn types when evaluated on June 20.

**Table 1.** Effect of different corn types on leaf litter decomposition rate and the contents of non-structural carbohydrate and total nitrogen in leaf litter.

| Leaf variable | Sampling time | Bt-506 | Zheng 58 | Zhengdan 958 | P-value |
| --- | --- | --- | --- | --- | --- |
| Decomposition rate (%) | Total | 26.7 ±1.6b | 25.0 ±2.0b | 31.5 ±1.9a | **0.002** |
|  | April 20 | 20.6 ±1.4a | 19.8 ±1.5a | 19.7 ±1.5a | 0.886 |
|  | May 20 | 22.7 ±1.2b | 21.1 ±1.5b | 28.7 ±1.6a | **0.003** |
|  | June 20 | 36.8 ±2.1b | 34.0 ±3.0b | 46.0 ±2.6a | **0.009** |
| Non-structural carbohydrate (mg/g) | Total | 60.5 ±1.3b | 61.1 ±1.4b | 63.8 ±1.0a | **0.015** |
|  | April 20 | 65.5 ±1.0b | 66.1 ±0.9b | 69.8 ±0.8a | **0.006** |
|  | May 20 | 62.8 ±0.8b | 64.0 ±0.6b | 67.7 ±1.0a | **0.001** |
|  | June 20 | 53.1 ±2.2a | 53.1 ±2.7a | 54.0 ±1.1a | 0.937 |
| Total nitrogen (mg/g) | Total | 2.0 ±0.1a | 2.0 ±0.1a | 1.7 ±0.1b | **< 0.001** |
|  | April 20 | 2.1 ±0.1a | 2.0 ±0.1a | 1.5 ±0.1b | **0.004** |
|  | May 20 | 2.0 ±0.1a | 2.1 ±0.1a | 1.8 ±0.1b | **0.018** |
|  | June 20 | 2.0 ±0.1a | 2.0 ±0.1a | 1.8 ±0.1a | 0.085 |

Each value represents mean ± SE. Different lowercases within a line indicate significantly different at $P < 0.05$ .

### The decomposition rate of corn leaf

Repeated measure ANOVA showed that the decomposition rate of leaf litters of different corns were significantly different (Table 1, *P* = 0.002), and that of the local type Zhengdan 958 was significantly higher than that of Bt-506 and Zheng 58 types. One-way ANOVA analysis showed that Bt-506 and Zheng 58 types had similar leaf litter decomposition rate for each of the three sampling dates, while Zhengdan 958 was significantly higher in the decomposition rate of leaf litter than Bt-506 and Zheng 58 types when sampled on May 20 and June 20, but the three corn types did not show significant difference for the samples on April 20 (Table 1).

### Collembolan abundance and Shannon-Wiener index

A total of 14,707 collembolans, involved in 18 species, were extracted from all the three corn litterbags in 2014. Among them, the dominant species was *Proisotoma minuta*, accounting for 93.7% of the total collembolan number. In addition, *Allonychiurus songi, Desoria* sp2 and *Folsomia bisetosa* were common species, accounting for 1.6, 2.4, and 1.2% respectively, while the number of other collembolan species was very few. There were 1,864, 1,970 and 10,873 collembolans in Bt-506, Zheng 58 and Zhengdan 958 litterbags, respectively (Table 2). The collembolan community from each of the three corn litterbags was also dominated by *P. minuta*.

**Table 2.** The number and percentage of different Collembola species in litterbags of different corn types.

| Collembola | Bt-506 |  | Zheng 58 |  | Zhengdan 958 |  | Total |  |
| --- | --- | --- | --- | --- | --- | --- | --- | --- |
|  | Number | Percentage (%) | Number | Percentage (%) | Number | Percentage (%) | Number | Percentage (%) |
| *Allonychiurus songi* | 78 | 4.18 | 89 | 4.52 | 69 | 0.63 | 236 | 1.60 |
| *Tullbergia yosii* | 2 | 0.11 | 0 | 0.00 | 1 | 0.01 | 3 | 0.02 |
| *Proisotoma minuta* | 1512 | 81.12 | 1709 | 86.75 | 10570 | 97.21 | 13791 | 93.77 |
| *Parisotoma notabilis* | 2 | 0.11 | 4 | 0.20 | 9 | 0.08 | 15 | 0.10 |
| *Desoria* sp1 | 0 | 0.00 | 1 | 0.05 | 0 | 0.00 | 1 | 0.01 |
| *Desoria* sp2 | 198 | 10.62 | 105 | 5.33 | 53 | 0.49 | 356 | 2.42 |
| *Folsomia bisetosa* | 10 | 0.54 | 21 | 1.07 | 139 | 1.28 | 170 | 1.16 |
| *Folsomides parvulus* | 2 | 0.11 | 2 | 0.10 | 1 | 0.01 | 5 | 0.03 |
| *Isotomedes* sp1 | 3 | 0.16 | 8 | 0.41 | 2 | 0.02 | 13 | 0.09 |
| *Isotomedes* sp2 | 12 | 0.64 | 5 | 0.25 | 7 | 0.06 | 24 | 0.16 |
| *Orchesellides sinensis* | 1 | 0.05 | 0 | 0.00 | 4 | 0.04 | 5 | 0.03 |
| *Entomobrya* sp1 | 10 | 0.54 | 13 | 0.66 | 5 | 0.05 | 28 | 0.19 |
| *Entomobrya koreana* | 30 | 1.61 | 11 | 0.56 | 7 | 0.06 | 48 | 0.33 |
| *Homidia phjongiangica* | 0 | 0.00 | 0 | 0.00 | 1 | 0.01 | 1 | 0.01 |
| *Orchesellides sinensis* | 1 | 0.05 | 0 | 0.00 | 5 | 0.05 | 6 | 0.04 |
| *Hypogastrura* sp1 | 2 | 0.11 | 1 | 0.05 | 0 | 0.00 | 3 | 0.02 |
| *Sminthuride* sp1 | 1 | 0.05 | 0 | 0.00 | 0 | 0.00 | 1 | 0.01 |
| *Sminthuride* sp2 | 0 | 0.00 | 1 | 0.05 | 0 | 0.00 | 1 | 0.01 |
| Total | 1864 | 100.00 | 1970 | 100.00 | 10873 | 100.00 | 14707 | 100.00 |

For the total collembolan samples collected at three times, repeated measure ANOVA showed that both the abundance of total collembolan species and the abundance of *P. minuta* did not show significantly different between Bt-506 and Zheng 58, they were both significantly lower than that in Zhengdan 958 (Table 3). As for the samples at each time, one-way ANOVA showed that the abundance of total collembolan species and *P. minuta* from the litterbags of Bt-506 were not significantly different from that of its near isoline Zheng 58, but they were both significantly lower than those from local type Zhengdan 958 when collembolans were sampled on May 20. The two indexes were not significantly different among the three corn types on April 20 and on June 20.

**Table 3.**
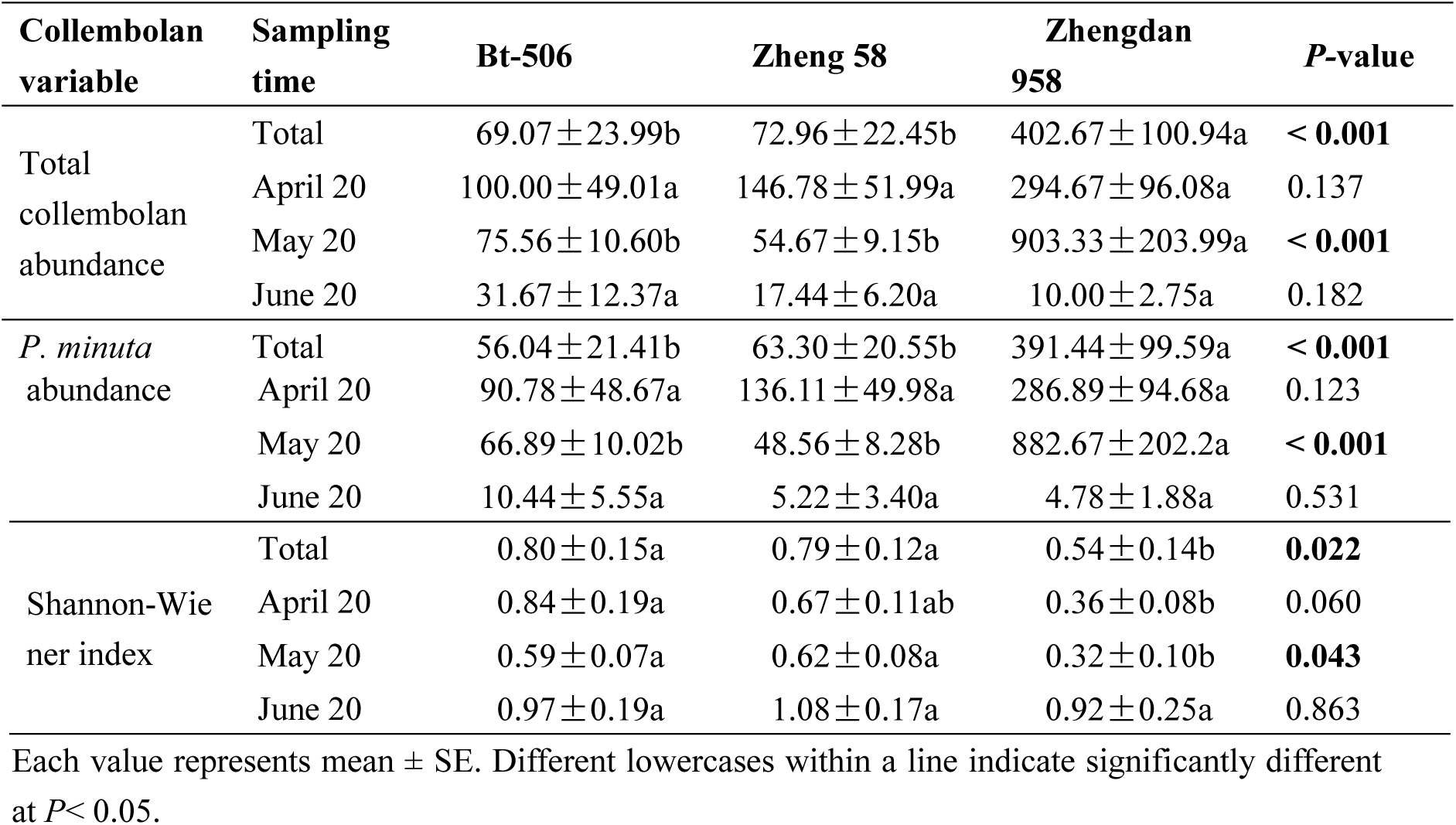
Effects of different corn types on abundance and Shannon-Wiener index of Collembola in litterbags.

The Shannon-Wiener index of the total collembolan species collected at three times from Bt-506 and Zheng 58 were not significantly different when analyzed by repeated measure ANOVA, while they were both significantly higher than that from Zhengdan 958 when analyzed by repeated measure ANOVA. The Shannon-Wiener indexes of the collembolan samples at each time from Bt-506 and Zheng 58 were also similar, which was revealed by one-way ANOVA, and they both had not significant difference compared to that from Zhengdan 958 on June 20. However the differences among the three corn types were not consistent at every time. For example, ANOVA showed that on April 20 the Shannon-Wiener index of the collembolan from Bt-506 was higher than that from Zheng 58 and Zhengdan 958, while it was nearly the same as that from Zheng 58 and higher than that from Zhengdan958 on May 20 (Table 3).

### The correlation among collembolan abundance, leaf litter decomposition rate, non-structural carbohydrate content and total nitrogen content

The collembolan abundance was significantly positively correlated with the non-structural carbohydrate content and decomposition rate of leaf litter, and was significantly negatively correlated with the total nitrogen content of leaf litter for the samples on May 20 (Table 4). The decomposition rate of leaf litter was also significantly positively correlated with its non-structural carbohydrate content, and was significantly negatively correlated with its total nitrogen content when we sampled on May 20, (Table 4). However, there was no significant correlation among collembolan abundance, leaf litter decomposition rate, non-structural carbohydrate content and total nitrogen content when sampling on 20 April or on 20 June (Table 4).

**Table 4.** Correlation coefficient between collembolan abundance, leaf litter decomposition rate and leaf litter indexes (including non-structural carbohydrate content, total nitrogen content).

|  | Sampling time | Non-structural carbohydrate content | Total nitrogen content | Decomposition rate |
| --- | --- | --- | --- | --- |
| Collembolan abundance | April 20 | 0.379 | -0.111 | 0.370 |
|  | May 20 | **0.412\*** | **-0.544\*\*** | **0.683\*\*** |
|  | June 20 | 0.269 | 0.105 | 0.121 |
| Decomposition rate | April 20 | -0.144 | -0.312 |  |
|  | May 20 | **0.549\*\*** | **-0.536\*\*** |  |
|  | June 20 | 0.130 | -0.323 |  |

\* and \*\* mean significantly correlated at 0.05 and 0.01.

## Discussion

Our study clearly showed that the Bt protein content of Bt-506 residue was decreased over time. The degradation of Bt protein is relatively slow in the first five months due to the low temperature, after April, it was accelerated significantly with the temperature increasing in Northeast China, but 8.1% still remained after buried into field for 7 months. However, there were no significant differences in leaf litter decomposition rate and collembolan abundance between Bt-506 and its near isoline Zheng 58 types in all sampling times. These results were the same as most of the previous studies on Bt corn. For example, Zurbrügg et al. [6] and Xiao et al. [41] have found that the decomposition rates were not different between Bt and non-Bt crops but among transgenic hybrids and among conventional hybrids through litterbag studies. Grigioni et al. [42], Zwahlen et al. [9] and Hönemann et al. [5] also found that the litters of Bt corns did not affect collembolan community. Besides, the Bt litters from other Bt crops, such as Bt cotton and Bt rice [43], did not affect collembolan community either. These results indicated that the Bt protein of Bt-506 event did not affect the decomposition rate of Bt-506 leaf litter and the collembolan community in the field in short term.

However, repeated measures ANOVA showed that the decomposition rates of leaf litter, the abundances of total Collembola species and the abundance of the dominant species *P. minuta* in the little bag of local type Zhengdan 958 were all significantly higher than that of Bt-506 and its near isoline Zheng 58 types. This might be due to the significant different chemical compositions of different corn leaves.

In this study, we found that the local type Zhengdan 958 had higher non-structural carbohydrate content and lower total nitrogen content than Bt-506 and its near isoline Zheng 58 on April 20 and May 20. Some previous studies found that the total nitrogen amount of litter influenced litter decomposition rate and colonization by fungi and bacteria in microcosms [13, 14]. And most of the soil Collembola was feed on these microorganisms [18], for example the dominant species *P. minuta* in our study is just a kind of fungivore [44], this species may be very abundant in the agriculture soil due to its preference for particular kinds of organic material [45]. At the same time, our study also showed that the abundance of collembolan and the litter decomposition rate of leaf litter were significantly correlated with the non-structural carbohydrate and total nitrogen contents of leaf litter, and the collembolan abundance was significantly positively correlated with leaf litter decomposition rate on May 20. Thus the higher decomposition rate of leaf litter and more abundance of collembolan for local type Zhengdan 958 were probably caused by the higher non-structural carbohydrate content and lower total nitrogen content in local corn Zhengdan 958.

The decomposition rate of leaf litter was not significant different among the three corn types, and there was no significant correlation among collembolan abundance, leaf litter decomposition rate, and the contents of non-structural carbohydrate and contents of total nitrogen on April 20, this may be due to the low temperature at that time. In Northeast China, as the temperature is very low in experimental field from November to April (most of the time is below 0 °C), when the soil fauna are in dormancy stage or inactive, the decomposition of litter residue was restricted.

With temperature rising, the soil fauna began to become active, and the leaf residue began to be decomposed quickly. By June 20, nearly half of the local type Zhengdan 958 residue was decomposed, and about a third of litters of Bt-506 and its near isoline Zheng 58 were decomposed. With leaf litter reducing, the collembolan abundance and the non-structural carbohydrate and total nitrogen contents of leaf litter in litterbags became similar for the three corn types; and there was no more significant correlation among collembolan abundance, leaf litter decomposition rate, contents of non-structural carbohydrate and contents of total nitrogen at this time.

## Conclusions

Our field study revealed that in the black soil region of China, Bt protein from transgenic corn plant didn’t significantly influenced the decomposition rate of leaf litter and the abundance of Collembola in Bt-506 litter bag when compared with its near isoline Zheng 58 in short term. Local type possesses higher non-structural carbohydrate content but lower total nitrogen content compared with Bt-506 and its near isoline Zheng 58 types, which probably was the main reason causing a higher decomposition rate of leaf litter and a higher abundance of Collembola in the leaf litterbags of local type Zhengdan 958.

## Supporting information

### S1 File. Raw data

The collembolan composition, leaf litter decomposition rate and the non-structural carbohydrate content and total nitrogen content in the leaf litters were included.

## Acknowledgements

We thank Professor Louis Deharveng from the Muséum national d’Histoire naturelle, France for identifying some of Collembola species. We thank Professor Jinsheng Lai from China Agricultural University for providing all corn seeds. We also thank Professor Sina Adl from University of Saskatchewan, Canada for revising the language. This project was supported by the Genetically Modified Organisms Breeding Major Projects, China (2016ZX08011-003), the National Nature Science Fund of China (31500345), and the innovation project of Jilin Academy of Agriculture Sciences, China (C6215000222).

## Author Contributions

### Conceptualization

Baifeng Wang, Xinyuan Song.

### Supervision

Xinyuan Song.

### Investigation

Baifeng Wang, Fengci Wu, Junqi Yin.

### Data analysis

Baifeng Wang, Fengci Wu, Ling Zhang, Xinyuan Song.

### Writing - original draft

Baifeng Wang, Fengci Wu, Xinyuan Song.

### Writing -review & editing

Baifeng Wang, Fengci Wu, Xinyuan Song.

## References

1. Shelton AM, Zhao JZ, Roush RT. Economic, ecological, food safety, and social consequences of the deployment of Bt transgenic plants. Ann Rev Entomol. 2002; 47: 845–881. https://doi.10.1146/annurev.ento.47.091201.145309.

2. Glaser JA, Matten SR. Sustainability of insect resistance management strategies for transgenic Bt corn. Biotechnol Adv. 2003; 22: 45–69. https://doi.10.1016/j.biotechadv.2003.08.016.

3. Hurley TM, Langrock I, Ostlie K. Estimating the benefits of Bt corn and cost of insect resistance management exante. J Agr Resour Econ. 2006; 31: 355–375. https://doi.10.1016/j.japwor.2004.12.005.

4. Zwahlen C, Hilbeck A, Gugerli P, Nentwig W. Degradation of the Cry1Ab protein within transgenic bacillus thuringiensis corn tissue in the field. Mol Ecol. 2003; 12: 765–775. https://doi.10.1046/j.1365-294X.2003.01767.x.

5. Hönemann L, Zurbrügg C, Nentwig W. Effects of Bt-corn decomposition on the composition of the soil meso- and macrofauna. Appl Soil Ecol. 2008; 40: 203–209. https://doi.10.1016/j.apsoil.2008.04.006.

6. Zurbrügg C, Hönemann L, Meissle M, Romeis J, Nentwig W. Decomposition dynamics and structural plant components of genetically modified Bt maize leaves do not differ from leaves of conventional hybrids. Transgenic Res. 2010; 19: 257–267. https://doi.10.1007/s11248-009-9304-x.

7. Kamota A, Muchaonyerwa P, Mnkeni PNS. Decomposition of surface-applied and soil-incorporated Bt maize leaf litter and Cry1Ab protein during winter fallow in South Africa. Pedosphere. 2014; 24: 251–257. https://doi.10.1016/S1002-0160(14)60011-4.

8. Böttger R, Schaller J, Lintow S, Gert Dudel E. Aquatic degradation of Cry1Ab protein and decomposition dynamics of transgenic corn leaves under controlled conditions. Ecotox Environ Safe. 2015; 113: 454–459. https://doi.10.1016/j.ecoenv.2014.12.034.

9. Zwahlen C, Hilbeck A, Nentwig W. Field decomposition of transgenic Bt maize residue and the impact on non-target soil invertebrates. Plant Soil. 2007; 300: 245–257. https://doi.10.1007/s11104-007-9410-6.

10. Böttger R, Schaller J, Lintow S, Gert Dudel E. Aquatic degradation of Cry1Ab protein and decomposition dynamics of transgenic corn leaves under controlled conditions. Ecotox Environ Safe. 2015; 113, 454–459. https://doi.10.1016/j.ecoenv.2014.12.034.

11. Saxena D, Stotzky G. Bt corn has a higher lignin content than non-Bt corn. Am J Bot. 2001; 88: 1704–1706. https://doi.10.2307/3558416.

12. Fang M, Motavalli PP, Kremer RJ, Nelson KA. Assessing changes in soil microbial communities and carbon mineralization in Bt and non-Bt corn residue-amended soils. App Soil Ecol. 2007; 37: 150–160. https://doi.10.1016/j.apsoil.2007.06.001.

13. Henriksen TM, Breland TA. Nitrogen availability effects on carbon mineralization, fungal and bacterial growth, and enzyme activities during decomposition of wheat straw in soil. Soil Biol Biochem. 1999; 31: 1121–1134. https://doi.10.1016/S0038-0717(99)00030-9.

14. Güsewell S, Gessner MO. N: P ratios influence litter decomposition and colonization by fungi and bacteria in microcosms. Funct Ecol. 2010; 23: 211–219. https://doi.10.1111/j.1365-2435.2008.01478.x.

15. Escher N, Käch B, Nentwig W. Decomposition of transgenic Bacillus thuringiensis maize by microorganisms and woodlice Porcellio scaber (Crustacea: Isopoda). Basic Appl Ecol. 2000; 1: 161–169. https://doi.10.1078/1439-1791-00024.

16. Icoz I, Stotzky G. Fate and effects of insect-resistant Bt crops in soil ecosystems. Soil Biol Biochem. 2008; 40: 559–586. https://doi.10.1016/j.soilbio.2007.11.002.

17. Hopkin SP. Biology of the Springtails (Insecta: Collembola)[M]//Biology of the springtails (Insecta, Collembola). Oxford University Press. 1997.

18. Rusek J. Biodiversity of collembola and their functional role in the ecosystem. Biodivers Conserv. 1998; 7: 1207 – 1219. https://doi.10.1023/a:1008887817883.

19. Bardgett RD, Putten WHVD. Belowground biodiversity and ecosystem functioning. Nature. 2014; 515: 505–511. https://doi.10.1038/nature13855.

20. Rebek EJ, Hogg DB, Young DK. Effect of four cropping systems on the abundance and diversity of epedaphic Springtails (Hexapoda: Parainsecta: Collembola) in Southern Wisconsin. Environ Entomol. 2002; 31: 37–46. https://doi.10.1603/0046-225X-31.1.37.

21. Santorufo L, Cortet J, Arena C, Goudon R, Rakoto A, Morel JL, et al. An assessment of the influence of the urban environment on collembolan communities in soils using taxonomy and trait-based approaches. Appl Soil Ecol. 2014; 78: 48–56. https://doi.10.1016/j.apsoil.2014.02.008.

22. Rossetti I, Bagella S, Cappai C, Caria MC, Lai R, Roggero PP, et al. Isolated cork oak trees affect soil properties and biodiversity in a mediterranean wooded grassland. Agric Ecosyst Environ. 2015; 202: 203–216. https://doi.10.1016/j.agee.2015.01.008.

23. Moore JC, Berlow EL, Coleman DC, Ruiter PC, Dong Q, Hastings A, Johnson NC, McCann KS, Melville K, et al. Detritus, trophic dynamics and biodiversity. Ecol Lett. 2004; 7: 584–600. https://doi.10.1111/j.1461-0248.2004.00606.x.

24. Endlweber K, Ruess L, Scheu S. Collembola switch diet in the presence of plant roots thereby functioning as herbivores. Soil Biol Biochem. 2009; 41: 1151–1154. https://doi.10.1016/j.soilbio.2009.02.022.

25. Bitzer RJ, Rice ME, Pilcher CD, Pilcher CL, Lam WF. Biodiversity and community structure of epedaphic and euedaphic springtails (Collembola) in transgenic rootworm Bt corn. Environ Entomol. 2005; 34: 1346–1376. https://doi.10.1603/0046-225X(2005)034[1346:BACSOE]2.0.CO;2.

26. Chang L, Liu XH, Ge F. Effect of elevated O3 associated with Bt cotton on the abundance, diversity and community structure of soil Collembola. Appl Soil Ecol. 2011; 47: 45–50. https://doi.10.1016/j.apsoil.2010.10.013.

27. Arias-Martní M, Garcaí M, Luciáñez MJ, Ortego F, Castañera P, Farinós GP. Effects of three-year cultivation of Cry1Ab-expressing Bt maize on soil microarthropod communities. Agr Ecosyst Environ. 2016; 220: 125–134. https://doi.10.1016/j.agee.2015.09.007.

28. Song XY, Chang L, Reddy Gadi VP, Zhang L, Fan CM, Wang BF. Use of taxonomic and trait-based approaches to evaluate the effects of transgenic Cry1Ac corn on the community characteristics of soil Collembola. Environ Entomo. 2019; 48: 263-269. Environ Entomo 48: 263–269. https://doi.10.1093/ee/nvy187.

29. Bellinger PF, Christiansen KA, Janssens F. Checklist of the Collembola of the World. http://www.collembola.org. (accessed 14 May 2019); 2019.

30. Heckmann LH, Griffiths BS, Caul S, Thompson J, Pusztai-Carey M, Moar WJ, Andersen MN, Krogh PH. Consequences for *Protaphorura armata* (Collembola: Onychiuridae) following exposure to genetically modified Bacillus thuringiensis (Bt) maize and non-Bt maize. Environ Pollut. 2006; 142: 212–216. https://doi.10.1016/j.envpol.2005.10.008.

31. Yuan YY, Ke X, Chen FJ, Krogh PH, Ge F. Decrease in catalase activity of *Folsomia candida* fed a Bt rice diet. Environ Pollut. 2011; 159: 3714–3720. https://doi.10.1016/j.envpol.2011.07.015.

32. Yuan YY, Xiao NW, Krogh PH, Chen FJ, Ge F. Laboratory assessment of the impacts of transgenic Bt rice on the ecological fitness of the soil non-target arthropod, *Folsomia candida* (Collembola: Isotomidae). Transgenic Res. 2013; 22: 791–803. https://doi.10.1007/s11248-013-9687-6.

33. SzabóB, Seres A, Bakonyi G, SzabóB, Seres A, Bakonyi G. Long-term consumption and food replacement of near-isogenic by Bt-maize alter life-history traits of *Folsomia candida* willem 1902 (Collembola). Appli Ecol Env Res. 2017; 15: 1275–1286. https://doi.10.15666/aeer/1504_12751286.

34. Zhang B, Yang Y, Zhou X, Shen P, Peng Y, Li Y. A laboratory assessment of the potential effect of cry1ab/cry2aj-containing Bt maize pollen on *Folsomia candida* by toxicological and biochemical analyses. Environ Pollut. 2017; 222: 94–100. https://doi.10.1016/j.envpol.2016.12.079.

35. Macfadyen A. Improved funnel-type extractors for soil arthropods. J. Anim Ecol. 1961; 30: 171–184. https://doi.10.2307/2120.

36. Christiansen K, Bellinger P. The Collembola of North America north of the Rio Grand. Grinnell College, Grinnell, Iowa; 1980.

37. Pomorski RJ. Onychiurinae of Poland (Collembola: Onychiuridae). Polish Taxonomical Society, Wrocklaw; 1998.

38. Yin WY. Pictorial Keys to Soil Animals of China. Science Press, Beijing, pp. 282-292, 592-600; 1998.

39. Potapov M. Synopses on Palaearctic Collembola: Isotomidae. Abhandlungen und Berichte des Naturkundemuseums Görlitz; 2001.

40. Tissue DT, Wright SJ. Effects of seasonal water availability on phenology and the annual shoot carbohydrate cycle of tropical forest shrubs. Funct Ecol. 1995; 9: 518–527. https://doi.10.2307/2390018.

41. Xiao MQ, Fang CM, Dong SS, Tang X, Chen Y, Yang SM, et al. Litterbag decomposition of litters from *Bacillus thuringiensis* (Bt) rice hybrids and the parental lines under multiple field conditions. J. Soils Sediment. 2014; 14: 1669–1682. https://doi.10.1007/s11368-014-0933-1.

42. Grigioni M, Daniele C, D’Avenio G, Barbaro V. Field decomposition of transgenic Bt maize residue and the impact on non-target soil invertebrates. Plant Soil. 2007; 300: 245–257. https://doi.10.1007/s11104-007-9410-6.

43. Bai YY, Yan RH, Ye GY, Huang FN, Cheng JA. Effects of transgenic rice expressing Bacillus thuringiensis Cry1Ab protein on ground-dwelling collembolan community in postharvest seasons. Environ. Entomol. 2010; 39: 243–251. https://doi.10.1603/EN09149.

44. Harasymek L, Sinha RN. Survival of Springtails *Hypogastrura tullbergi* and *Proisotoma minuta* on Fungal and Bacterial diets. Environ Entomol. 1974; 3: 965–968. https://doi.10.1093/ee/3.6.965.

45. Santorufo L, Cortet J, Nahmani J, Pernin C, Salmon S, Pernot A, et al. Responses of functional and taxonomic collembolan community structure to site management in Mediterranean urban and surrounding area. Eur J Soil Biol. 2015; 70: 46–57. https://doi.10.1016/j.ejsobi.2015.07.003.

